# A practical staging atlas to study embryonic development of *Octopus vulgaris* under controlled laboratory conditions

**DOI:** 10.1101/2020.01.13.903922

**Authors:** Deryckere Astrid, Styfhals Ruth, Vidal Erica A.G., Almansa Eduardo, Seuntjens Eve

## Abstract

**Background:** *Octopus vulgaris* has been an iconic cephalopod species for neurobiology research as well as for cephalopod aquaculture. It is one of the most intelligent and well-studied invertebrates, possessing both long- and short-term memory and the striking ability to perform complex cognitive tasks. Nevertheless, how the common octopus developed these uncommon features remains enigmatic. *O. vulgaris* females spawn thousands of small eggs and remain with their clutch during their entire development, cleaning, venting and protecting the eggs. In fact, eggs incubated without females usually do not develop normally, mainly due to biological contamination (fungi, bacteria, etc.). This high level of parental care might have hampered laboratory research on the embryonic development of this intriguing cephalopod.

**Results:** Here, we present a completely parameter-controlled artificial seawater standalone egg incubation system that replaces maternal care and allows successful embryonic development of a small-egged octopus species until hatching in a laboratory environment. We also provide a practical and detailed 1 staging atlas based on bright-field and light sheet fluorescence microscopy imaging for precise monitoring of embryonic development. The atlas has a comparative section to benchmark stages to the different scales published by Naef (1928), Arnold (1965) and Boletzky (2016). Finally, we provide methods to monitor health and wellbeing of embryos during organogenesis.

**Conclusion:** Besides introducing the study of *O. vulgaris* embryonic development to a wider community, this work can be a high-quality reference for comparative evolutionary developmental biology.

## Background

*Octopus vulgaris* is a marine carnivorous cephalopod mollusk that inhabits a variety of coastal areas in a wide distributional range (1). Almost a century ago, Naef published the first classification of the embryonic development of *Loligo vulgaris, Sepia officinalis, O. vulgaris*, and *Argonauta argo*, demonstrating their potential of becoming model systems in developmental biology (2).

Cephalopod eggs can be roughly divided in small, medium or large in size and show a great diversification of encapsulation mechanisms (3). While the common cuttlefish lays individual medium-sized encapsulated eggs covered by an ink stained multilayer gelatinous envelope, the common octopus produces small eggs with a single transparent chorionic coat, devoid of a protective gelatinous capsule, which significantly increases their ease of use in laboratory experimental studies. The chorion itself is drawn out into a stalk and in octopods, many stalks are interwoven and glued together with material secreted by the female oviducal glands to form a string or festoon (Fig. 1A) (4, 5). Octopuses that lay eggs that hatch out as planktonic paralarvae generally produce thousands of small eggs, reaching 500,000 in *O. vulgaris* (6). Fertilization is achieved during spawning whereafter the string is attached to a substrate in the den (3, 6). During embryonic development, cephalopod eggs generally increase in volume, although this phenomenon is more pronounced in decabrachian eggs compared to octopod eggs (7). In *O. vulgaris* eggs, this swelling process affects egg width and wet weight whereas length is nearly unaffected (8).

**Figure 1.**
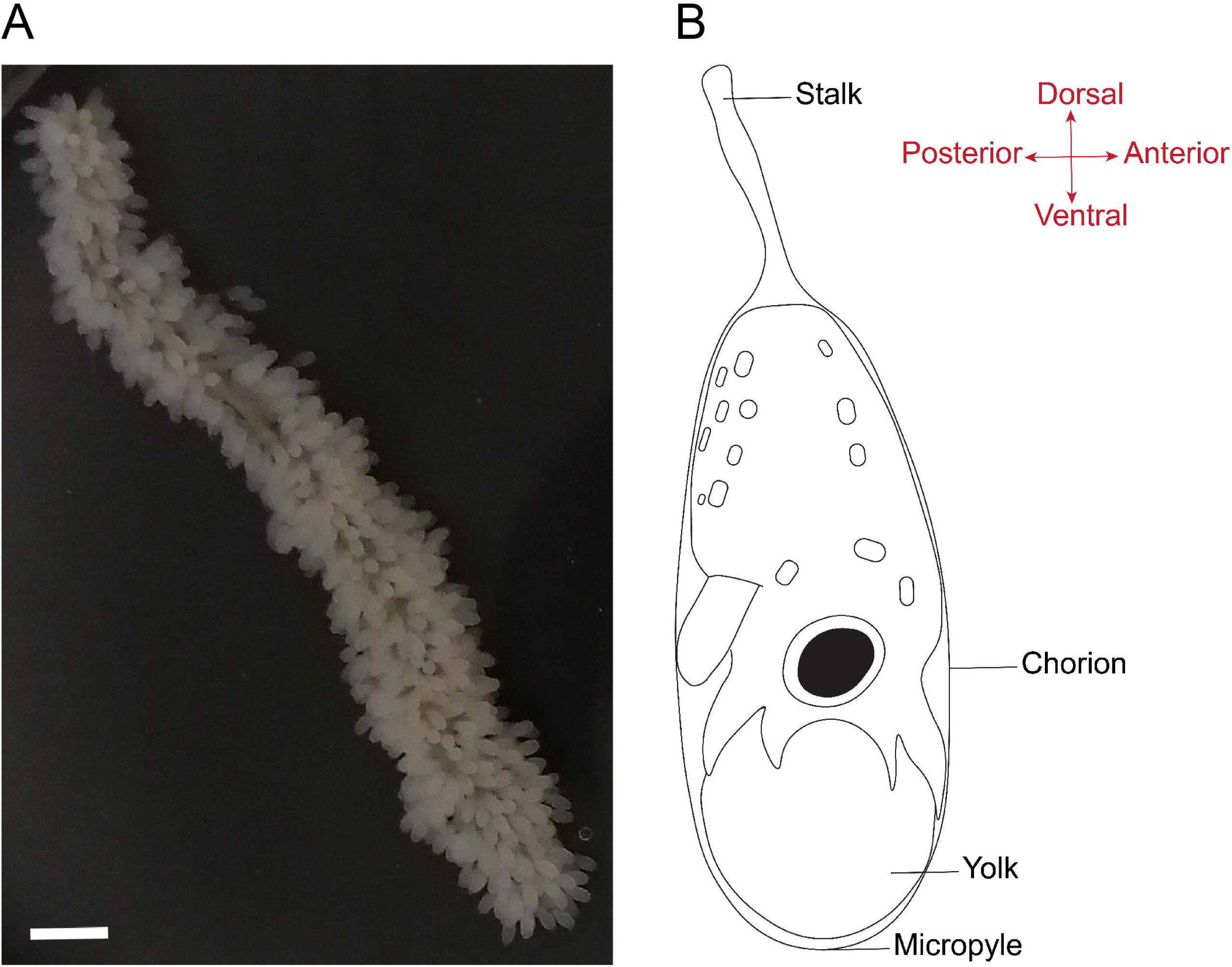
*Octopus vulgaris* eggs and the embryonic morphological body axes. A. A string of *O. vulgaris* eggs. Scale bar represents 500 μm. B. The morphological axes in cephalopod embryos correspond to the axes in other mollusks. In this orientation, the location of the funnel is posterior, the embryonic mouth is anterior, the arms are ventral and the mantle is dorsal.

The embryonic development of cephalopods can roughly be separated in three periods. The first one includes maturation and fertilization of the oocyte, discoidal meroblastic cleavage to form the blastodisc and division to complete the blastoderm, the latter annotated as Stage I by Naef. The gastrulation or second period comprises the formation of the germinal layers with establishment of endoderm and extra-embryonic yolk epithelium (Stage II-IV) and the start of epiboly followed by concentrations of mesoderm (Stage V-VII). The organogenesis or third period begins with an elevation of blastodisc folds that prelude the appearance of the first organ primordia (Stage VIII-XI) that will give rise to the typical dibranchiate topology (Stage XII-XVII) and then, linear growth will eventually form a fully developed hatchling (Stage XVIII-XX) (2). The last stages of development (maturation) are more difficult to compare between cephalopods, since species that produce large eggs generally hatch out as juveniles that are miniature adults, while small egg-embryos hatch out as small planktonic paralarvae. The latter still have to go through major morphological changes to attain the juvenile form, such as the development of the arm-crown complex, swimming control, the chromatophore system and horizontal pupillary response (9–11). Furthermore, taxon specific features that arise in cuttlefish (e.g. cuttlebone) or squid (e.g. tentacles) embryos are absent from octopus and thus not discussed here.

Octopuses (e.g. *Octopus, Eledone* and *Tremoctopus*) undergo double reversion during embryonic development (12, 13). The first reversion or blastokinesis takes place at Stage VII in *O. vulgaris*, when the extra-embryonic yolk epithelium just completed closure at the vegetative pole and is realized by a change of direction of the ciliary beat of the yolk envelope (12). In this process, the embryo migrates from the micropyle to the stalk side of the egg, which takes 7 to 36 hrs depending on water temperature (12, 14). While positioning at the stalk side might protect embryos better from predators and would reduce mechanical stress during organogenesis (Nande, personal communication), failure of turning does not impact embryonic development. The second reversion at Stage XIX then positions the embryo for smooth hatching (12). The physiological and morphological factors that trigger hatching in cephalopods are still unknown (5, 15), but hatching starts with stretching mantle movements that rupture the apex of cells in the hatching gland or organ of Hoyle at the dorsal tip on the mantle (16, 17). These glandular cells store proteolytic enzymes that dissolve the chorion locally, making the egg integument permeable to water, which increases the osmotic pressure within the perivitelline space (5,18–20). Afterwards, the mantle is extruded due to a release of pressure and the Kölliker organs (hard bristle-like structures spread over the skin) make sure that the embryo does not slip back into the chorion so it can move freely from the egg during hatching (9,20,21).

Due to breeding season limitations as well as geographical spread, different cephalopod species are being researched around the world. In addition, the release of several cephalopod genomes as well as transcriptomic information over the last years now allows molecular and functional studies on these enigmatic creatures (22–26). In combination with novel genome editing technologies, this opens interesting opportunities to interrogate *in vivo* gene function. However, in *O. vulgaris*, progress in these fields has been hampered by the absence of protocols to maintain egg clutches without maternal care in standardized laboratory conditions. Furthermore, to fully evaluate the impact of genetic change on development, an updated description of embryonic development using modern imaging technologies is valuable. Additionally, there is a need for a standardized, fully-illustrated staging system allowing easy comparison of embryonic development between different cephalopod model species. We acknowledge the inevitable generalization introduced by comparing embryonic stages and refer to species-specific morphological descriptions of *S. officinalis, Euprymna scolopes, Todarodes pacificus, Loligo pealei, L. gahi* and *O. vulgaris* (2,27–31). Although cephalopod egg and thus hatchling size and consequently the embryonic development duration greatly vary, morphogenetic processes are similar.

Naefs staging atlas of *O. vulgaris* is still frequently used today, although being based on the age of embryos in days (which changes according to incubation temperature (8)), rather than stage-specific morphological characteristics. Therefore, Arnold and Lemaire (later adapted by Boletzky) introduced ten extra stages, focusing on the development of *L. pealei* (officially renamed *Doryteuthis pealeii*) and *S. officinalis*, respectively (27,28,32). These extra stages mostly cover the period of embryo cleavage (e.g. Arnold and Boletzky Stage 9 correspond to Naef Stage I), which can also be described by the number of blastomeres as proposed by Naef. Both staging scales do not readily cover the considerable gaps in development between different stages and important developmental events are still largely neglected. Moreover, the arbitrary use of 20 or 30 stage atlases by different research groups make evolutionary comparison between cephalopods challenging. Therefore, we introduce here a comprehensive atlas, based on the staging presented by Naef, but including a differentiation in early and late phases of some stages in an attempt to highlight important details and to cover larger developmental gaps. We focus on the development of *O. vulgaris* as a model for small-egged cephalopod species. Furthermore, for easy translation between cephalopod species, we propose a comparative table of staging scales most often used by cephalopod researchers, ie. Arnold stages for *D. pealei*, Boletzky stages for *S. officinalis* and Naef stages for *O. vulgaris* (2,27,28). Hence, this atlas not only represents a timely standardized staging system to allow easy comparison between different model species, but also provides accompanying images to easily illustrate important developmental features. Detailed bright-field and light sheet fluorescence microscopy (LSFM) images of all developmental stages were added to be used in the laboratory as a staging atlas.

Finally, this work describes a standardized standalone tank system that should facilitate any laboratory on small-egged cephalopods, regardless of access to fresh seawater. We also supply validation assays for checking the health of embryos at different stages.

## Results

The small, yolky eggs of *O. vulgaris* are roughly 2.5 mm long and 1 mm wide. Octopus embryos are described to develop poorly without maternal care (2, 33). However, we have found that *O. vulgaris* embryos can develop without maternal care in artificial oxygenated seawater at continuous strong flow rate and dim light. The standalone system ensured a continuous flow in the tanks resulting in an oblique orientation and soft swirling of the strings, likely mimicking the jet flow the mother normally provides (Fig. 2). The embryos developed highly synchronous within the string and hatched after approximately one month at 19 °C. We provide a summary table with key characteristics of each stage to allow consistent staging of *O. vulgaris* embryos (Table 1) as well as a comparative table including Arnold and Boletzky stages for easy translation between cephalopods (Table 2). As the developmental stages presented by Naef are based on days of development rather than on morphological characteristics and contain considerable gaps in development, we split some events and added ‘.1’ or ‘.2’ in such cases. For all descriptions presented, the morphological axes of the embryo are used (Fig. 1B). According to these axes, the location of the funnel is posterior, the embryonic mouth anterior, the arm crown ventral and the mantle dorsal.

**Figure 2.**
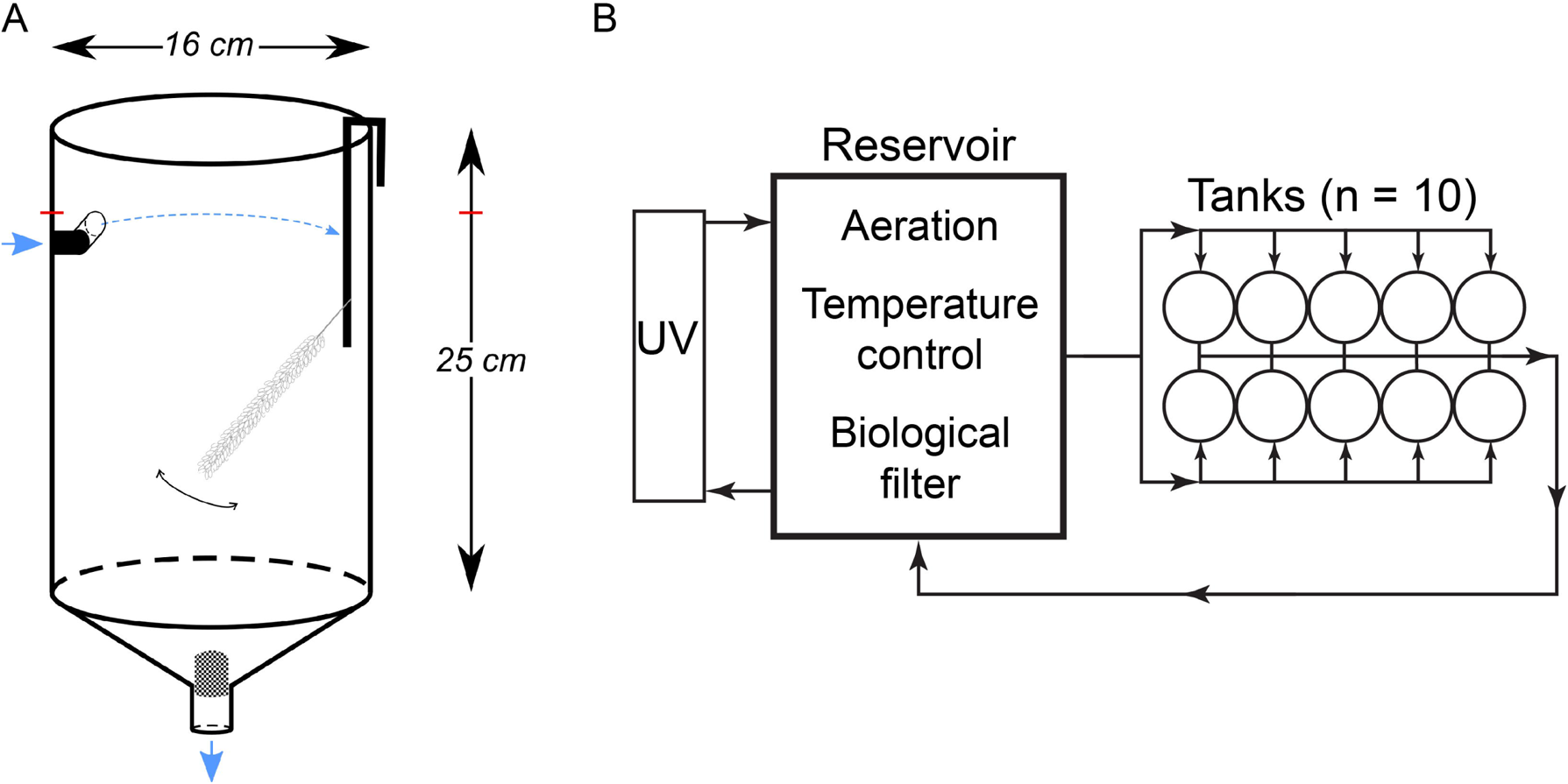
Graphical representation of the standalone system used for egg incubation. A. The opaque conocylindrical PVC tank has a water inlet at the top (blue arrow), which creates a circular current and delivers seawater at an exchange rate of 3 L min^-1^. A mechanical filter (1 mm mesh size) is placed at the bottom of the tank in the water outflow. An *O. vulgaris* egg string is attached to a glass rod and placed on the lateral side of the tank, where it moves constantly by a gentle current generated by the water inflow. The blue arrows indicate the water flow (from top to bottom) and the red horizontal bars indicate the water level. B. The standalone system consists of 10 conocylindrical tanks placed on top of a reservoir. Artificial seawater is aerated by the strong water flow pouring into the biological filter in the reservoir and is sterilized by an external UV filter (details provided in Methods section). The total volume in the system is 100 L.

### Cleavage, Gastrulation and Epiboly

The germinal disc is restricted to the animal pole of the egg, at the micropyle side, which is opposite from the stalk. Meroblastic, bilaterally symmetrical cleavage and subsequent formation of the blastodisc (Stage I) takes place over the first 24-48 hrs after fertilization, depending on water temperature. The first three cleavages are incomplete and generate eight equally sized blastomeres in octopods (Fig. 3A-D), which differs from decapods where the two dorso-medial cells are more narrow compared to the ventro-medial cells (2). Further cell proliferation results in the formation of the blastodisc at Stage I (Fig. 3E). At Stage II, formation of the blastula is completed (Fig. 3F), followed by the onset of epiboly at Stage III, characterized by lateral expansion of the blastoderm over the yolk by cell division (Fig. 3G). The blastodisc, which can be found at the very top of the yolk at Stage II starts to grow and expand over the yolk, generating a cap-like structure by Stage IV (Fig. 3H). At Stage V, a quarter of the yolk is covered by the embryonic cap (Fig. 3I). Using bright-field imaging, the embryo looks uniform at this stage. However, using light sheet microscopy and DAPI as a nuclear stain, the embryo proper with its densely packed nuclei can be clearly distinguished from the extraembryonic ectoderm with larger nuclei spaced further apart (Fig. 3I-I’). At Stage VI, the germinal disc covers half of the yolk mass (Fig. 3J-J’). From this stage onwards, the embryo slowly rotates clockwise when observed from the micropyle side of the egg, along its longitudinal axis (Additional file 1 shows a movie of embryo rotation accelerated to 8x original speed at Stage XI) (12, 14). By the end of Stage VII.1, the embryo and yolk envelope (extraembryonic) cover 3/4 th of the yolk, followed by complete closure at the vegetative pole, ready for the first reversion (Fig. 3K).

**Figure 3.**
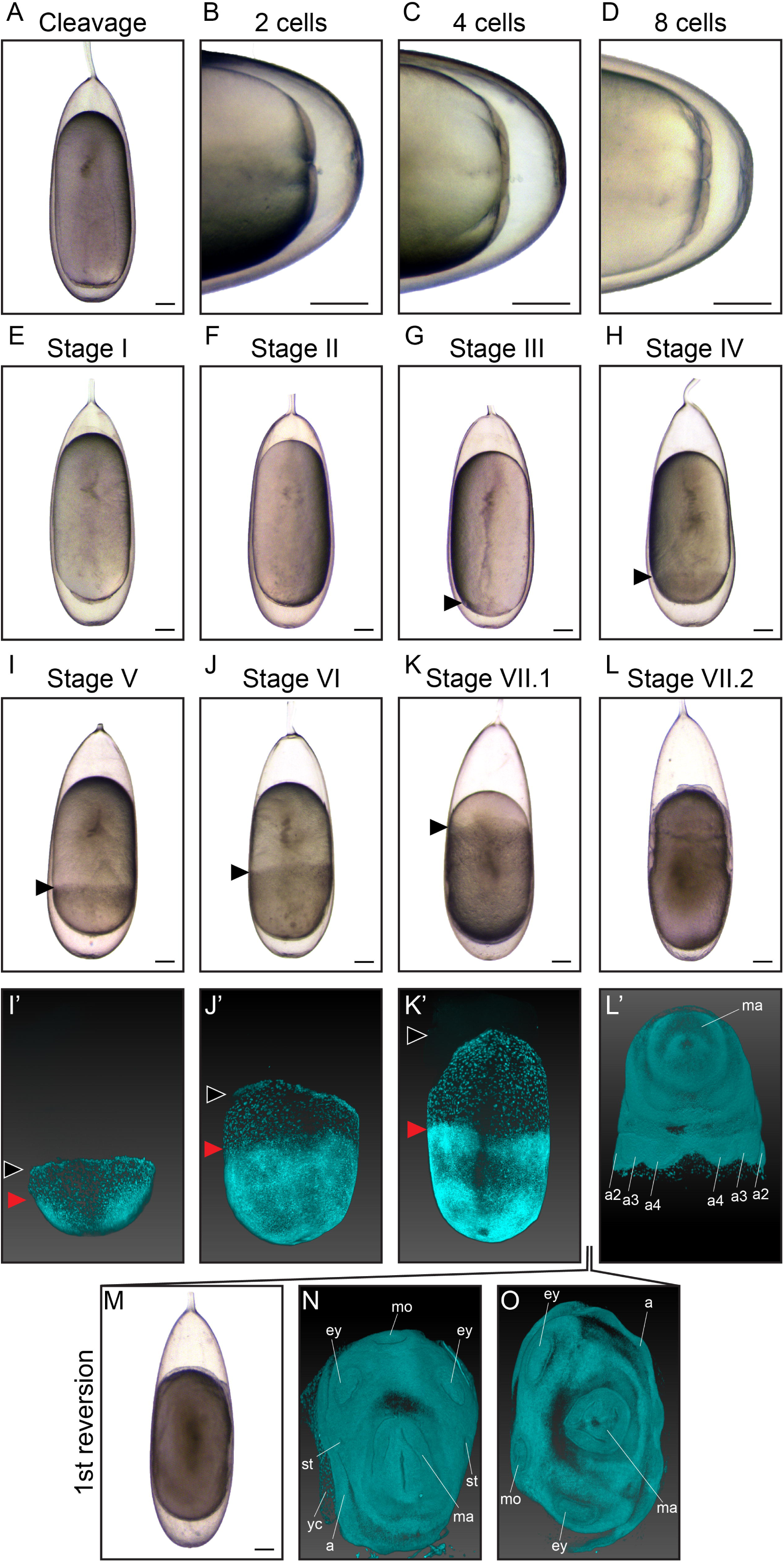
Cleavage, gastrulation, epiboly and reversion in *O. vulgaris*. Bright-field images of embryos in cleavage (A-D), at Stage I (E), Stage II (F), Stage III (G), Stage IV (H), Stage V (I), Stage VI (J), Stage VII.1 (K) and Stage VII.2 (L). Nuclear staining of Stage V (I’), Stage VI (J’), Stage VII.1 (K’) and Stage VII.2 (L’) embryos imaged with LSFM. Black arrowheads indicate the progression of epiboly, red arrowheads the borders of the embryo proper. At Stage VII, *O. vulgaris* embryos undergo the first reversion (M) and can be observed in distinct phases/ topologies during reversion (N-O) with LSFM. Scale bars represent 200 μm. *Abbreviations: a, arm; ey, eye; LSFM, light sheet fluorescence microscopy; ma, mantle; mo, mouth; st, statocyst; yc, yolk cells*.

### Organogenesis and Maturation

At Stage VII.1, the surface of the embryo appears smooth. The first organ primordia can be visualized using DAPI, revealing the prospective arms as patches of dense nuclei close to the yolk envelope (Fig. 3K-K’). The embryo makes its first reversion at the end of Stage VII. This process takes 7 to 36 hrs, depending on the incubation temperature (14), in which the embryo migrates over the yolk from the micropyle to the stalk side of the egg and can be observed in different topologies (Fig. 3M-O). At Stage VII.2, primordia become visible by bright-field microscopy as thickenings and depressions that arise from the surface of the embryo (Fig. 3L). The eye placodes, mantle anlage, arm primordia and mouth are the first distinguishable structures (Fig. 3L’) and become more discernable towards Stage VIII (Fig. 4), when the mantle rim is elevated.

**Figure 4.**
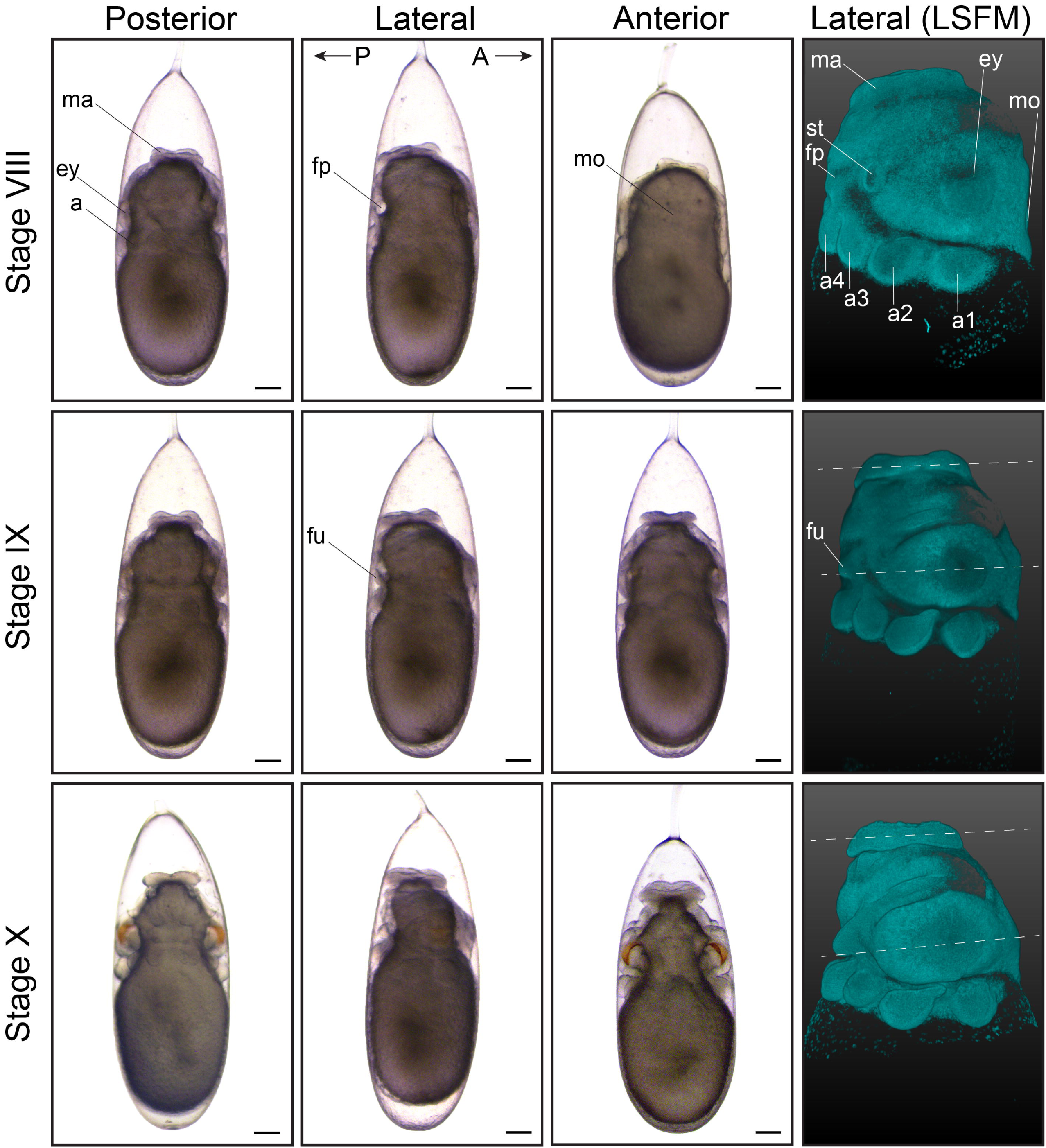
First part of organogenesis in *O. vulgaris*. Bright-field images of *O. vulgaris* embryos from Stage VIII to Stage X from the posterior, lateral and anterior side. Lateral LSFM images after DAPI staining show that the planes that run through the mantle and eyes run parallel (white dashed lines). Scale bars represent 200 μm. *Abbreviations: A, anterior; a, arm; D, dorsal; ey, eye; fp, funnel pouch; fu, funnel; LSFM, light sheet fluorescence microscopy; ma, mantle; mo, mouth; P, posterior; st, statocyst; V, ventral*.

**Figure 5.**
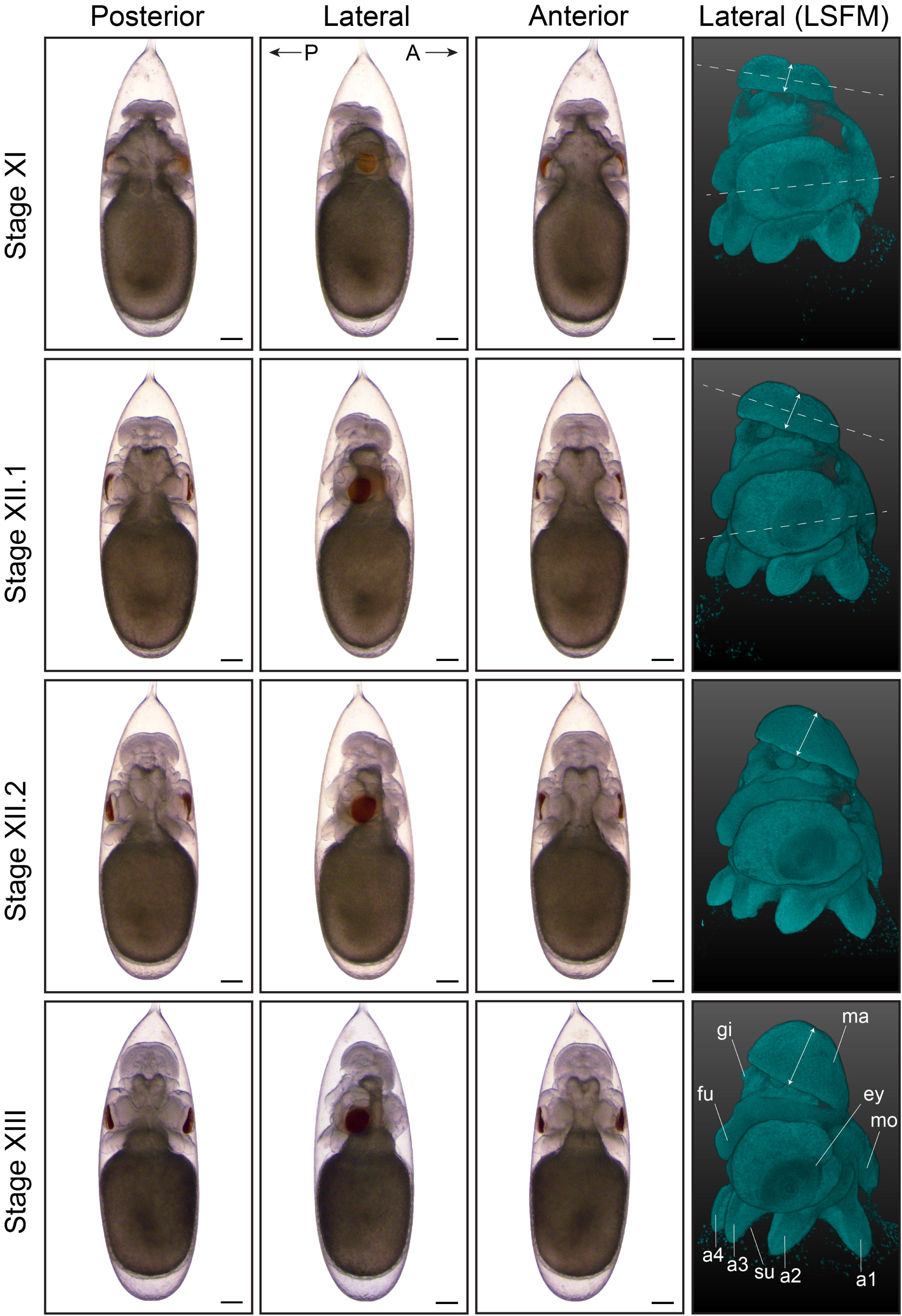
Second part of organogenesis in *O. vulgaris*. Bright-field images of *O. vulgaris* embryos from Stage XI to Stage XIII from the posterior, lateral and anterior side. Lateral LSFM images after DAPI staining show how the mantle is now tilted (white dashed lines) and growing (white double arrows) during development. Scale bars represent 200 μm. *Abbreviations: A, anterior; a, arm; D, dorsal; ey, eye; fu, funnel; gi, gills; LSFM, light sheet fluorescence microscopy; ma, mantle; mo, mouth; P, posterior; st, statocyst; su, sucker; V, ventral*.

During the next stages of organogenesis, the organ primordia become more prominent and are clearly distinguishable from the yolk, giving rise to an immature embryo at Stage XVII (Fig. 4-7; Additional files 2-13 show movies of embryos imaged with LSFM). At Stage IX, the arm buds are clearly separated from one another, the mantle appears more elevated and first yellow pigmentation of the retina is visible. The yolk sac envelope that contains blood lacuna and a network of muscular elements starts to create peristaltic waves of surface contraction at this stage, establishing blood circulation for the early embryo (Additional file 14 shows yolk contraction at Stage XI) (34). This phenomenon will cease around Stage XVI, when the embryonic heartbeat is well established and when the area of contact between the yolk envelope and the chorion becomes too small (12).

In order to distinguish embryos between Stages IX and XIII, mantle size and the angle relative to the imaginary plane through the eyes, as well as folding of the funnel tube are easily recognizable morphological characteristics (LSFM images in Fig. 4, 5, funnel in Fig. 6). The shape of the funnel is visible through the chorion, but is easier to observe after dechorionation. At Stage IX, the funnel tube rudiments become visible (Fig. 6A) and fuse at the margins by stage X (white arrow Fig. 6B). At Stage XI, the funnel tube rudiments have grown in size and bend towards the midline (Fig. 6C). Then, at the beginning of Stage XII (Stage XII.1), the funnel starts to form a real tube that is fused at the ventral extremity by Stage XII.2 (Fig. 6D-E). But, it is at Stage XIII that the formation of the siphon shaped funnel tube is complete (Fig. 6F). In the subsequent events, the position of the mouth on the anterior side changes (Fig. 7, white arrows on LSFM images). The mouth is situated between the first pair of arms on the anterior side from Stage VIII to XIV and is still open to the outside at Stage XV.1. It will start to internalize, becoming encircled by the anterior arms at Stage XV.2. By Stage XVI, the mouth is covered by the arm crown, waiting to take its final position as soon as the outer yolk is reduced.

**Figure 6.**
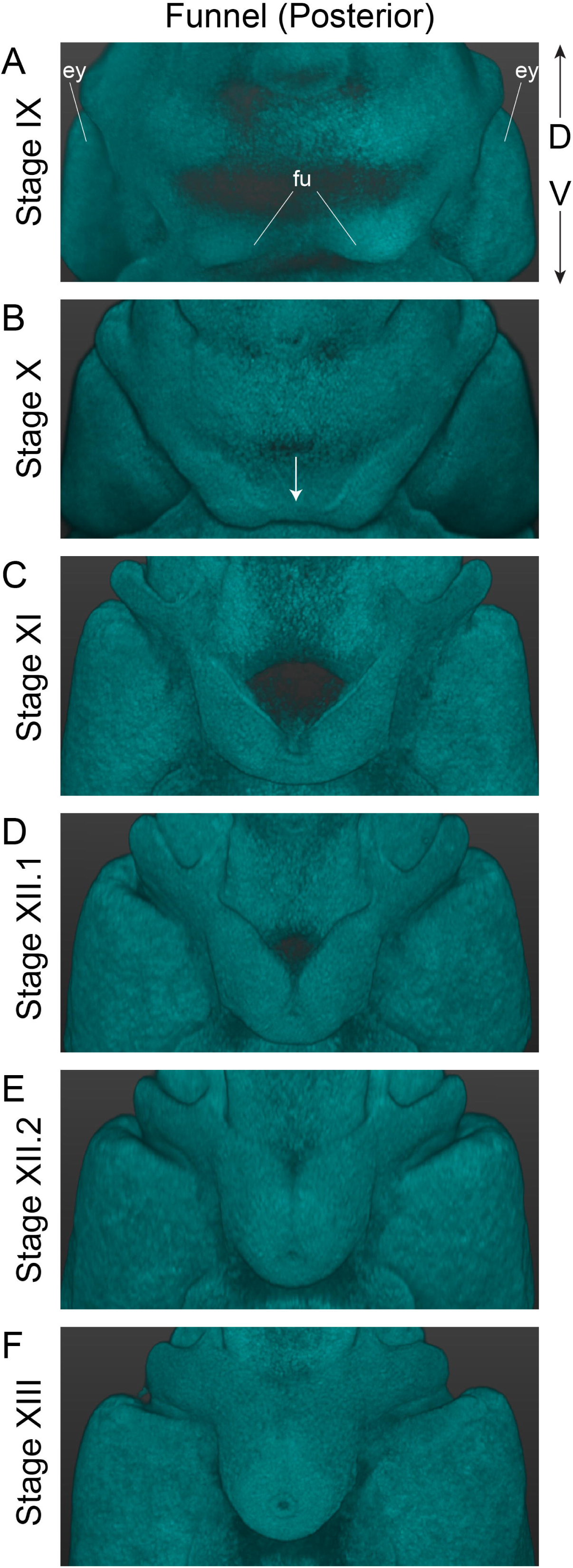
Development of the funnel apparatus in *O. vulgaris*. LSFM image of the posterior side of the embryo focusing on the funnel apparatus, showing its gradual thickening and fusion to form a funnel tube by Stage XIII. The funnel rudiments visible at Stage IX (A) fuse at the ventral margins at Stage X (white arrow in B). The rudiments then bend towards the midline at Stage XI (C) until they are touching one another at Stage XII.1 (D). Fusion to form the tube starts at the ventral extremity at Stage XII.2 (E) and closure finishes at the dorsal side by Stage XIII (F). *Abbreviations: D, dorsal; ey, eye; fu, funnel; LSFM, light sheet fluorescence microscopy; V, ventral*.

**Figure 7.**
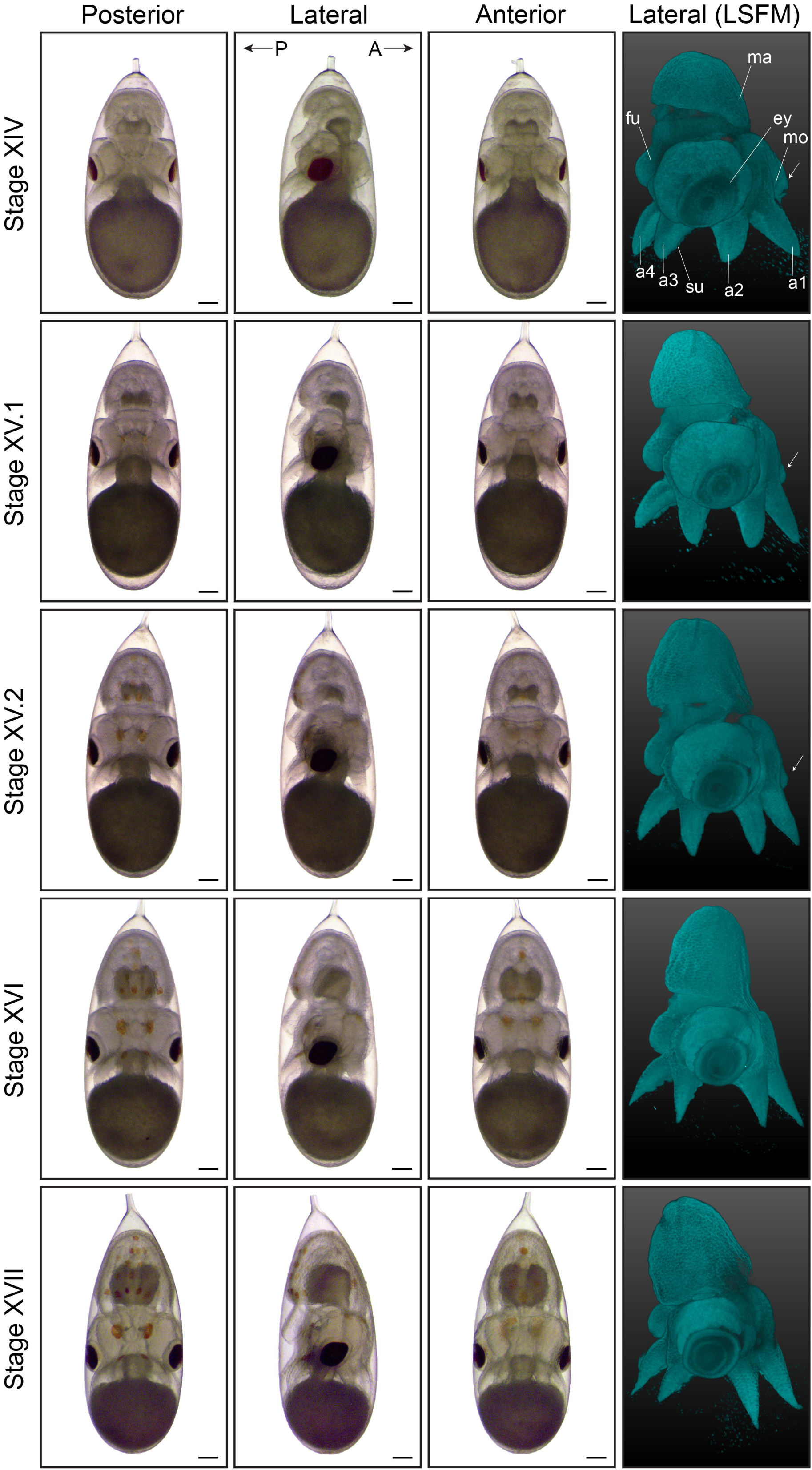
Third part of organogenesis in *O. vulgaris*. Bright-field images of *O. vulgaris* embryos from Stage XIV to Stage XVII from the posterior, lateral and anterior side. The appearance of chromatophores on the posterior and subsequently anterior side can be used to stage the embryos. Lateral LSFM images after DAPI staining show the internalization of the mouth (white arrows) with the mouth lying outside the arm crown at Stage XIV and inside by Stage XVI. Scale bars represent 200 μm. *Abbreviations: A, anterior; a, arm; ey, eye; fu, funnel; gi, gills; LSFM, light sheet fluorescence microscopy; ma, mantle; mo, mouth; P, posterior; su, sucker*.

**Figure 8.**
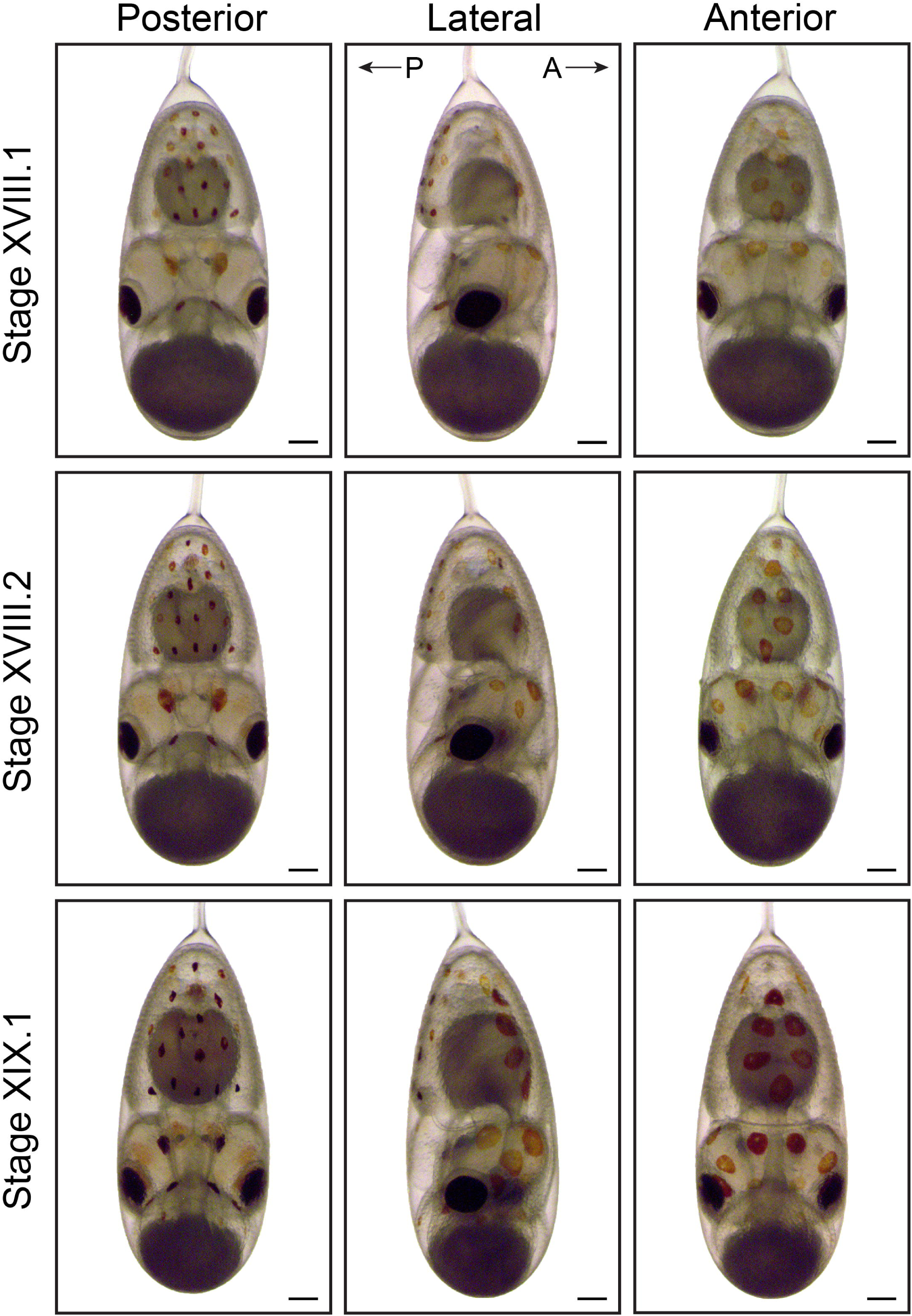
Final stages of maturation in *O. vulgaris* (Part 1) Bright-field images of *O. vulgaris* embryos from Stage XVIII.1 to XIX.1 from the posterior, lateral and anterior side. The chromatophore pattern (number, size and color) and the size of the external yolk sack can be used to distinguish the different stages before hatching. Scale bars represent 200 μm.

**Figure 9.**
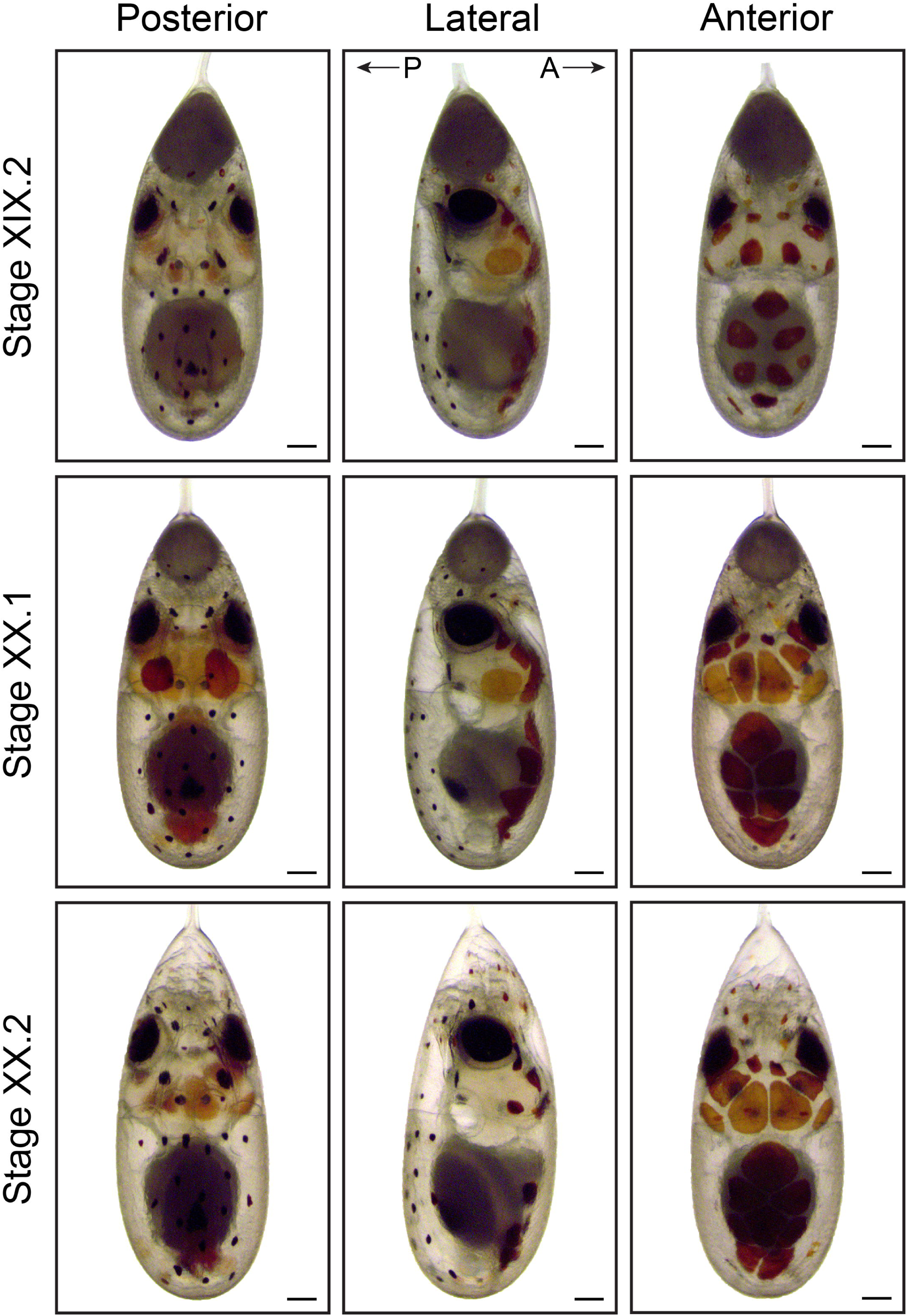
Final stages of maturation in *O. vulgaris* (Part 2) Bright-field images of *O. vulgaris* embryos from Stage XIX.2 to XX.2 from the posterior, lateral and anterior side. The chromatophore pattern (number, size and color) and the size of the external yolk sack can be used to distinguish the different stages before hatching. After the second reversion between Stage XIX.1 and XIX.2, (premature) hatching can occur. Scale bars represent 200 μm.

As the embryos grow, the shape of the mantle goes from depressed towards the middle at Stage VII.2 to flat and perpendicular to the longitudinal axis at Stage X. At Stage XI, the mantle is elevated on the posterior side and thus tilted and clearly grows in size by Stage XII. At Stages XIII and XIV, the length of the mantle equals and exceeds the length of the head in the dorsoventral axis, respectively (Fig. 5, 7), and at Stage XIV, a heartbeat can be observed at the mantle tip (Additional file 15 shows embryonic heart beat at Stage XVII). From Stage IX to stage XIV, the color of the retina changes from light yellow to dark red/brown. The color of the eye and retina continues to darken during development, until the eye is completely black and covered by an iridescent layer, clearly visible from Stage XIX onwards.

The chromatophore pattern (appearance, color and size of chromatophores) is another convenient characteristic to stage *O. vulgaris* embryos (Fig. 7, 8, 9). At Stages XV.1 and XV.2, the first chromatophores appear as small yellow dots on the posterior side, next to the funnel and on the mantle, respectively. By Stage XVI, the first chromatophores on the anterior mantle appear. From Stage XVIII.2 onwards, the chromatophores react to changes in light intensity under the microscope (expand under light stimulation and contract in the dark). The ratio of the size of the external yolk sack in relation to the size of the embryo is another measure that can be used for staging (Fig. 7, 8, 9). At Stage XIV, this ratio approximates 1:1 and rapidly decreases to 1:3 at Stage XVI, 1:4 at Stage XVIII.1 to 1:6 at Stage XIX.1. This latter stage is also characterized by the first appearance of ink in the ink sac on the posterior side. The embryo undergoes the second reversion at Stage XIX. We annotate these stages as XIX.1 before and XIX.2 after the second reversion.

At Stages XX.1 and XX.2, the external yolk sack is nearly and completely depleted, respectively (Additional file 16 shows a movie of a Stage XX.2 embryo imaged with LSFM). It has been described that cephalopod embryos are likely slightly sedated in the egg by a tranquillizing factor to prevent premature hatching which can occur at these stages (35). What precisely induces natural hatching is still unknown, but it is easily triggered by several factors, such as mechanical stimuli, photoperiodicity and sudden changes in light levels or temperature (15). We observed that natural hatching starts approximately seven days after the second reversion at 19 °C, but is detrimental to the paralarvae in the tank system under continuous flow. Therefore, seven days post second reversion, we moved the strings from the system to a different tank containing aerated artificial seawater, which induced hatching within minutes.

**Table 1.** Hallmarks of each developmental stage in *O. vulgaris*.

| Stage | Characteristic |
| --- | --- |
| Stage 0 | Cleavage |
| Stage I | Morula (advanced cleavage) |
| Stage II | Blastula; disk flatter |
| Stage III | Onset of epiboly |
| Stage IV | Formation of the germinal disk, as wide as the yolk |
| Stage V | Epiboly reaches 1/4th of the yolk |
| Stage VI | Epiboly reaches 1/2nd of the yolk |
| Stage VII.1 | Epiboly reaches 3/4th of the yolk |
|  | Visible thickening of placodes starts |
| Stage VII.2 | Embryo completed first reversion |
|  | Primordia of eyes, mouth, mantle and arms clearly visible |
| Stage VIII | Mouth and eye invagination |
|  | Mantle elevated and embryo thicker |
|  | Funnel pouches visible in lateral view |
| Stage IX | First eye pigmentation (yellowish) |
|  | Primordia more prominent |
|  | Funnel tube rudiments are distinct |
|  | Contraction of yolk envelope evident |
| Stage X | Mantle flat, no depression in the middle |
|  | Eye vesicles sticking out with light orange retina ("saddle" shape) |
|  | Funnel tube rudiments fuse at the ventral margins |
| Stage XI | Mantle tilted |
|  | Funnel tube rudiments bended towards midline |
|  | Arm buds 'elevated' from yolk |
| Stage XII.1 | Mantle thicker and covers 1/2nd of gills |
|  | Funnel tube rudiments start to form a tube ventrally |
|  | First suckers recognizable on the posterior side |
| Stage XII.2 | Mantle bowl-shaped |
|  | Funnel formed siphon at ventral extremity |
| Stage XIII | Mantle is bigger |
|  | Formation of the funnel complete |
|  | Arms elongated and pointed with (3) prominent suckers |
| Stage XIV | Mantle as wide as long and cube-shaped, covers gills completely |
|  | Heartbeat starts |
|  | Embryo and yolk have equal size |
| Stage XV.1 | Mantle completely covers ventral margin of funnel |
|  | Inner yolk strongly constricted (connection inner and outer yolk sac very thin) |
|  | Appearance of two chromatophores laterally from the funnel |
| Stage XV.2 | Mouth encircled by anterior arms |
|  | First chromatophores on posterior mantle appear |
| Stage XVI | Mouth completely covered by arm crown |
|  | Yolk size 1/3rd of embryo + yolk |
|  | Few chromatophores on anterior mantle evident |
| Stage XVII | Chromatophores darker and more numerous |
| Stage XVIII.1 | Yolk size 1/4th of embryo + yolk |
|  | Posterior chromatophores darker and chromatophores appear next to the eye |
|  | Embryo more active in egg (mantle contraction) |
| Stage XVIII.2 | Anterior chromatophores darker. Chromatophores react to light stimulus |
| Stage XIX.1 | Eyes tilted and covered with iridophores |
|  | Yolk size 1/6th of embryo + yolk |
|  | Embryos react to mechanical stimulus |
|  | Pigmentation of ink sac |
| Stage XIX.2 | Embryos completed second reversion |
| Stage XX.1 | Minimal outer yolk sack |
|  | Chromatophore expansion and contraction more widely distributed |
| Stage XX.2 | Absence of outer yolk |
|  | Hatching |

### Assays to evaluate embryonic fitness

Yolk contraction can be observed from Stage IX to Stage XVI under the stereomicroscope and is a valuable readout to evaluate embryonic survival at early organogenesis stages. Furthermore, upon development of the retina, a “saddle” to discoidal shape of the pigmented layer is typical of high-quality embryos. Frowning or folding of the retina points towards poor health. From Stage XIV onwards, a heartbeat can be recognized in the transparent embryos. Occasionally, small crustaceans can be observed on the strings. Generally, these are part of the natural ecosystem of the string and are not impacting embryonic development. Nevertheless, poor rearing conditions (insufficient flow, dissolved oxygen levels and strings floating or sunken) can trigger strings to overgrow with fungi (white thread-like structures or parts turning pale or pink) or get infected by worms. A final readout of state of the art rearing is the hatching of actively swimming paralarvae that display positive phototaxis, reported for most cephalopod hatchlings (9,36,37).

**Table 2.** Comparative developmental guide for cephalopod development.

| *Octopus vulgaris* | *Doryteuthis pealeii* | *Sepia officinalis* | General characteristics | Stage |
| --- | --- | --- | --- | --- |
| **Adapted from Naef, 1928** | **Arnold, 1965** | **Boletzky, 2016** |  |  |
| Fertilized egg | Stage 1 | Stage 1 | Newly laid fertilized egg that did not finish maturation | Segmentation (cleavage) |
| First maturation division | Stage 2 | Stage 1 | First polar body | Segmentation (cleavage) |
| Second maturation division | Stage 3 | Stage 1 | 3 polar bodies | Segmentation (cleavage) |
| 2 cells, first cleavage | Stage 4 | Stage 2 | Two partially separated cells | Segmentation (cleavage) |
| 4 cells, second cleavage | Stage 5 | Stage 3 | Four partially separated cells | Segmentation (cleavage) |
| 8 cells, third | Stage 6 | Stage 4 | Eight incompletely separated | Segmentation |
| cleavage |  |  | cells | (cleavage) |
| 16 cells, fourth cleavage | Stage 7 | Stage 5 | 2 completely closed blastomeres, 14 blastocoques | Segmentation (cleavage) |
| 32 cells, fifth cleavage | Stage 8 | Stage 6 | 12 blastomeres, 20 blastocoques | Segmentation (cleavage) |
| 64-66 cells, sixth cleavage | Stage 9 | Stage 7 | 36 blastomeres, 28 blastocoques | Segmentation (cleavage) |
| Stage I | Stage 10 | Stage 8 | Morula (advanced cleavage) | Segmentation (cleavage) |
| Stage II | Stage 10 | Stage 9 | Blastula | Segmentation (cleavage) |
| Stage III | Stage 11 | Stage 10 | Onset of epiboly | Gastrulation |
| Stage IV | Stage 12 | Stage 11-12 | Formation of the germinal disk (yolk envelope & embryonic proper) | Gastrulation |
| Stage V | Stage 13 | Stage 13 | Epiboly continues | Gastrulation |
| Stage VI | Stage 14-15-16 | Stage 14 | Mesoderm concentrations become more distinct (smooth surface) | Epiboly |
| Stage VII.1 | Stage 17 | Stage 15 | Visible thickening of placodes starts | Epiboly |
| Stage VII.2 | Stage 18 | Stage 15 | Primordia of mantle, eyes, mouth and arms visible | Epiboly/<br>Organogenesis |
| Stage VIII | Stage 19 | Stage 16 | Mouth and eye invagination, funnel pouches and statocysts visible | Organogenesis |
| Stage IX | Stage 20 | Stage 17 | Primordia more prominent. Funnel tube rudiments are distinct | Organogenesis |
| Stage X | Stage 21 | Stage 18 | Eye vesicles closed and sticking out. Funnel tube rudiments fuse ventrally | Organogenesis |
| Stage XI | Stage 22 | Stage 19 | Mantle starts to grow. Funnel tube rudiments bend towards midline | Organogenesis |
| Stage XII.1 | Stage 23 | Stage 20 | Funnel tube rudiments start to form a tube ventrally. Mantle covers 1/2nd of gills | Organogenesis |
| Stage XII.2 | Stage 23 | Stage 20 | Funnel formed siphon at ventral extremity. Mantle bowl-shaped | Organogenesis |
| Stage XIII | Stage 23 | Stage 21 | Formation of the funnel complete. Iris fold rudiment visible | Organogenesis |
| Stage XIV | Stage 24 | Stage 22 | Mantle as wide as long and covers gills completely | Organogenesis |
| Stage XV.1 | Stage 25 | Stage 23 | Mantle covers ventral margin of funnel. Inner yolk strongly constricted | Organogenesis |
| Stage XV.2 | Stage 26 | Stage 24 | Mouth starts to internalize | Organogenesis |
| Stage XVI | Stage 27 | Stage 25-26 | Few anterior chromatophores evident. Mouth completely covered by arm crown | Organogenesis |
| Stage XVII | Stage 28 | Stage 27 | Mantle enlarged in relation to head. Chromatophores numerous | Organogenesis |
| Stage XVIII.1 | Stage 28 | Stage 27 | Yolk sac same size as head<br>(*Octopus & Loligo*) | Growth |
| Stage XVIII.2 | Stage 28 | Stage 27 | Chromatophores darker | Growth |
| Stage XIX.1-XIX.2 | Stage 29 | Stage 28 | Pigmentation of ink sac. Eyes<br>covered with iridophores | Growth |
| Stage XX.1 | Stage 29 | Stage 29 | Yolk nearly depleted | Growth |
| Stage XX.2 | Stage 30 | Stage 30 | Loss of outer yolk and hatchling | Growth |

## Discussion

We introduced a low-cost standalone system that runs on artificial seawater for incubating small-egged Octopus species without maternal care. The feasibility and effectiveness of our system was reflected in a highly synchronous development of embryos within the string and in the production of viable hatchlings.

### Replacing maternal care

Incirrate octopods and some oceanic squids display parental care during embryonic development (15, 38). As in many octopods, *O. vulgaris* females take care of the eggs during the whole embryonic development, venting, cleaning and protecting them from predators. Female care ensures high hatching rates and the production of viable hatchlings as incubating eggs without the female often resulted in the proliferation of pathogens (fungi and bacteria) on the eggs (EAG Vidal, personal observation)(39). Incubation without maternal care for small-egged Octopus species has therefore not always been possible. On the other hand, the large eggs (up to 17 mm length) of *Octopus maya* can be artificially incubated without the female with nearly 100% success rate for fertilized eggs (40). In 1977, Van Heukelem used a glass funnel with filtered seawater to incubate the eggs of *O. maya*. After adjusting seawater flow, the eggs were maintained slowly tumbling and rubbing against one another in order to keep the egg surface clean and aerated. This author also described that air bubbles interfered with the development of the yolk epithelium and were thus harmful to the embryos (41). Similarly, our early attempts to incubate egg strings in beakers or tanks with fine air bubbles venting in from the bottom were equally unsuccessful, and yielded embryos that did not manage to partition the inner from the outer yolk sack, leading to incomplete yolk epithelium development and thus, embryo malformation and death. Accordingly, egg strings should not be exposed to air bubbles and aeration of the water is therefore best performed outside of the tanks that house the strings. A second major improvement to our tank system was the combination of a relatively strong water flow and attachment of strings to the lateral side of the tank where the main current is, several centimeters below the water surface, ensuring that the strings were swirling around gently in the water. These adaptations yield a similar condition in which eggs are continuously rubbing against each other, likely functioning as a natural cleaning system. Third, we maintained the eggs in very dim light conditions (0-5 lux) using a 14L:10D photoperiod, which likely mimics the natural dark environment of egg clutches in the den. To what extent egg maintenance in dim light is absolutely required remains to be studied, but a preliminary study in *L. vulgaris* showed an inverse relationship between light intensity and hatching success (42).

### Hallmarks of good quality embryos

Using these conditions, we noted a highly synchronous development within each string, with very little embryonic death or malformation occurring. Whereas embryonic development progress is more difficult to assess before Stage VII.2, after the first reversion, a number of hallmarks can be used to assess vitality of the embryos, such as yolk contractions, and later on heart beating, although these might be irregular at early embryonic stages. Inability to gradually reduce the inner yolk during organogenesis, frowning of the retina and increased presence of particles on the chorion are signs of poor embryo condition, and resulted in embryonic death. Poor embryo condition also seemed to trigger an increased infestation risk of bacteria, fungi or parasites (worms). Recently, Maldonado et al. successfully used a bleaching protocol on *Octopus insularis* eggs to clean them from microorganism contamination prior to individual egg housing in restricted water circulation (43). Restricted housing without bleaching caused 100 % mortality within a few days whereas 67.6 % of the bleached embryos survived. Although individual egg housing can be beneficial for certain experiments, it is extremely labor intensive and requires much more space to house the same amount of eggs compared to the system described here.

### Developing clear staging criteria

Several hallmarks can be used to easily identify developmental stages in *O. vulgaris*. In the early embryo, the rate of epiboly demarcates each stage. Afterwards, from Stage IX to Stage XIV, the formation of the funnel, as well as mantle shape and size can be used to differentiate the embryos. From Stage XV onwards, the amount, color intensity and reactivity of the chromatophores increases with embryo development and the size of the outer yolk sack is progressively reduced until it is completely absorbed at hatching (Table 1). When rearing conditions are not ideal, premature hatching occurs and paralarvae hatch out with the outer yolk sack still present, resulting in high mortality rates (44).

When using these morphological characteristics to stage *O. vulgaris* embryos, we encountered the necessity to introduce extra developmental stages besides those proposed by Naef (2). In addition, these extra stages with defined hallmarks make the comparison with other cephalopods easier. For example, Stage VII annotated by Naef as the stage where differentiation of the mesoderm contractions starts, corresponds to Arnold Stages 17 and 18 in *L. pealei (D. pealeii)*. By dividing this Stage VII in two, Stage VII.1 now corresponds to Stage 17, where placode thickening starts and Stage VII.2 corresponds to Stage 18, where organ primordia of the mantle, eyes, mouth and arms are clearly visible (see also Table 2) (27).

In cephalopod research, two different representations of body axes are used at random (i.e. morphological and functional body axes). When adopting the morphological body axes of a cephalopod, the embryonic mouth is anterior and the funnel posterior, the mantle dorsal and the arms ventral. In this setup, the mouth-funnel axis corresponds to the molluscan anterior-posterior axis where the foot is ventral. On the other hand, when using the functional body axes that correspond to the adult convention, the embryonic mouth is dorsal and the funnel ventral, the mantle posterior and the arms anterior. For the sake of comparison, the body axes should be clearly defined in each publication.

## Conclusions

The data presented here aimed at facilitating developmental research on cephalopods, and in particular octopus species, under standardized laboratory conditions. We therefore removed potential roadblocks, such as obligatory maternal care and the availability of natural seawater, which we solved by introducing a low-cost standalone tank system that runs on artificial seawater. Given the high fecundity of *O. vulgaris* females, the high number of eggs from each string and the robustness of the embryos, egg strings from different females can be shipped and shared between laboratories in order to serve the growing community. In the present study, using classical and contemporary imaging technologies, we generated a comprehensive overview of *O. vulgaris* embryonic development along with a practical illustrated atlas. We documented the different stages of embryonic development and compared them to published literature, allowing practical use and unambiguous staging, which represents a reliable resource for comparative developmental biology in the cephalopod field.

## Methods

### Standalone system for egg incubation and embryo maintenance

Live egg strings of *O. vulgaris* were obtained from breeding females from the Instituto Español de Oceanografía (IEO), Tenerife, Spain, as soon as possible after spawning. The egg strings were attached to a nylon thread and transported in seawater in closed 50 mL falcons at ambient temperature to the Laboratory of Developmental Neurobiology in Leuven, Belgium. Transport time from tank to tank amounted to a maximum of 12 hrs. Upon arrival in the lab, single strings were placed in a standalone system that consisted of 10 conocylindrical opaque PVC tanks (16 cm diameter, 25 cm height), with a water inlet placed at the top to create a circular current with a water exchange rate of 3 L min^-1^ (Fig. 2A). The standalone system continuously circulated aerated artificial seawater (Instant Ocean 40 g L^-1^, supplemented with 8 mg L^-1^ Strontium), which was continuously cooled to 19 °C, sterilized by UV (Deltec Profi UV sterilizer 39 W type 391), filtered through a mesh (1 mm) in each tank and circulated through a shared biological filter (21 x 21 x 11 cm, MarinePure Block, CER MEDIA) (Fig. 2B). The total volume of the system was 100 L, conductivity 50-55 mS, light intensity between 0-5 lux (dusk-dark) with a photoperiod of 14L:10D and pH was maintained between 8.1-8.3.

Each *O. vulgaris* egg string was attached to a glass rod using the nylon threads and placed on the lateral side of a tank, where it was in constant motion generated by the gentle current from the water inflow (Fig. 2A). The top of the tanks was covered with plastic foil to avoid evaporation and Aluminum foil to block light. After observation of the second reversion, embryos were left undisturbed for seven days to avoid premature hatching (44).

### Bright-field imaging

Egg strings were obtained from four different females. Embryos were observed daily and a sample of 20 representative embryos was removed daily from the string for imaging. All observations were based upon embryos reared in the standalone system. At least 4 strings for each female were monitored. Since fertilization was not timed and spawning takes place over several days, different strings of a single female were in different developmental stages, allowing monitoring of subtle changes and transitions during embryonic development. Embryos reared in this system were compared to fixed reference embryos obtained from sibling strings at the laboratory of E. Almansa (IEO) and also to independent reference embryos from the laboratory of E. Vidal (Center for Marine Studies, University of Parana, Brazil). Images were taken with a Zeiss Stereo Discovery.V8 equipped with an AxioCam ICc 3 camera (Carl Zeiss AG, Germany) and represent static stages based on a morphological consensus from different embryos.

### Optimized CUBIC clearing protocol

The advanced CUBIC (Clear, Unobstructed Brain/Body Imaging Cocktails and Computational Analysis) protocol was adapted from Susaki et al. (45). In short, eggs were fixed overnight in 4% paraformaldehyde (PFA) in phosphate-buffered saline (PBS) – or from Stage XX.1 onwards first submersed in 2% EtOH in seawater (to avoid stress and premature hatching) and then fixed in 2% EtOH, 4% PFA in seawater - and washed in PBS. To anticipate retrieval and convenient manipulation of the cleared embryos, Chinese ink was injected in the yolk before manual dechorionation using forceps. Embryos were incubated in 1/2-destilled-water-diluted ScaleCUBIC-1 in an orbital shaker of a hybridization oven at 37 °C for 3-6 hrs and then immersed in ScaleCUBIC-1 (25 wt% urea, 25 wt% Quadrol, 15 wt% Triton X-100). After overnight incubation, ScaleCUBIC-1 was replaced and embryos were further incubated for three days with one additional ScaleCUBIC-1 replacement. At this point, the yolk was completely transparent, chromatophores were cleared and the eye pigment of Stage XX embryos was reduced from black to reddish (comparable to live Stage XIII embryos). Embryos were then washed with PBS three times (1x 2 h, 1x overnight and 1x 2h) in the hybridization oven. Afterwards, they were incubated in 1/2-water-diluted ScaleCUBIC-2 for 3-6 hrs (until the samples sunk to the bottom) and then incubated in ScaleCUBIC-2 (25 wt% urea, 50 wt% sucrose, 10 wt% triethanolamine) for one day in the hybridization oven. For nuclear staining, DAPI (final concentration 1 μg mL^-1^) was added to ScaleCUBIC-1 in the three days incubation in ScaleCUBIC-1 step and during washes in PBS.

### Light sheet fluorescence microscopy (LSFM)

Stained embryos were glued with their yolk sack on a metal rod and imaged using a Zeiss Z1 light sheet microscope (Carl Zeiss AG, Germany) in low-viscosity immersion oil mix (Mineral oil, Sigma M8410 and Silicon oil, Sigma 378488, 1:1). Then, 3D reconstructions were generated in Arivis (Vision4D, Zeiss Edition 2.10.5).

## Declarations

### Ethics approval and consent to participate

Not applicable

### Consent for publication

Not applicable

### Availability of data and materials

All data generated or analyzed during this study are included in this published article [and its supplementary information files].

### Competing interests

The authors declare that they have no competing interests

## Funding

A. Deryckere was supported by an SB Ph.D. fellowship of Fonds Wetenschappelijk Onderzoek (FWO) Belgium (Project No. 1S19517N) and R. Styfhals was supported by a fellowship from Stazione Zoologica Anton Dohrn and KU Leuven. E. Almansa was supported by the Spanish government (OCTOMICs project, AGL2017-89475-C2-2-R) and EAG. Vidal was supported by the Brazilian National Research Council-CNPq (Pro. 312331/2018-1 and 426797/2018-3). E. Seuntjens was supported by funding of the FWO (G0B5916N and G0C2618N) and the KU Leuven (C14/16/049). Publication cost was covered by the KU Leuven Fund for Fair Open Access.

### Authors’ contributions

EA provided live octopus eggs and EV provided fixed reference samples. AD and RS sampled octopus embryos. AD carried out the experiments. ES supervised the findings of this work. AD prepared the figures and wrote the manuscript with critical input from all authors. All authors read and approved the final manuscript.

## Acknowledgements

We wish to thank Beatriz C. Felipe and Mª Jesús Lago for their technical support in the broodstock management and Oliver Van Moerbeke and Arnold Van Den Eynde for designing and constructing the standalone system.

## Supplementary movies

### Additional file 1

.mov

Movie of embryo rotation at Stage XI

The embryo rotates around its longitudinal axis starting from Stage VI. This movie shows this movement accelerated to 8x the original speed.

### Additional file 2

.avi

Movie of a CUBIC cleared embryo stained with DAPI at Stage VIII

A Stage VIII CUBIC cleared, DAPI stained embryo imaged with LSFM shown rotating along the longitudinal axis.

### Additional file 3

.avi

Movie of a CUBIC cleared embryo stained with DAPI at Stage IX

A Stage IX CUBIC cleared, DAPI stained embryo imaged with LSFM shown rotating along the longitudinal axis.

### Additional file 4

.avi

Movie of a CUBIC cleared embryo stained with DAPI at Stage X

A Stage X CUBIC cleared, DAPI stained embryo imaged with LSFM shown rotating along the longitudinal axis.

### Additional file 5

.avi

Movie of a CUBIC cleared embryo stained with DAPI at Stage XI

A Stage XI CUBIC cleared, DAPI stained embryo imaged with LSFM shown rotating along the longitudinal axis.

### Additional file 6

.avi

Movie of a CUBIC cleared embryo stained with DAPI at Stage XII.1

A Stage XII.1 CUBIC cleared, DAPI stained embryo imaged with LSFM shown rotating along the longitudinal axis.

### Additional file 7

.avi

Movie of a CUBIC cleared embryo stained with DAPI at Stage XII.2

A Stage XII.2 CUBIC cleared, DAPI stained embryo imaged with LSFM shown rotating along the longitudinal axis.

### Additional file 8

.avi

Movie of a CUBIC cleared embryo stained with DAPI at Stage XIII

A Stage XIII CUBIC cleared, DAPI stained embryo imaged with LSFM shown rotating along the longitudinal axis.

### Additional file 9

.avi

Movie of a CUBIC cleared embryo stained with DAPI at Stage XIV

A Stage XIV CUBIC cleared, DAPI stained embryo imaged with LSFM shown rotating along the longitudinal axis.

### Additional file 10

.avi

Movie of a CUBIC cleared embryo stained with DAPI at Stage XV.1

A Stage XV.1 CUBIC cleared, DAPI stained embryo imaged with LSFM shown rotating along the longitudinal axis.

### Additional file 11

.avi

Movie of a CUBIC cleared embryo stained with DAPI at Stage XV.2

A Stage XV.2 CUBIC cleared, DAPI stained embryo imaged with LSFM shown rotating along the longitudinal axis.

### Additional file 12

.avi

Movie of a CUBIC cleared embryo stained with DAPI at Stage XVI

A Stage XVI CUBIC cleared, DAPI stained embryo imaged with LSFM shown rotating along the longitudinal axis.

### Additional file 13

.avi

Movie of a CUBIC cleared embryo stained with DAPI at Stage XVII

A Stage XVII CUBIC cleared, DAPI stained embryo imaged with LSFM shown rotating along the longitudinal axis.

### Additional file 14

.mov

Movie of external yolk contraction at Stage XI

The yolk sack of the embryo contracts between Stages IX and XVI and is shown here at Stage XI.

### Additional file 15

.avi

Movie of heart beat at Stage XVII

Heart beat can be observed from Stage XIV onwards and is shown here at Stage XVII.

### Additional file 16

.avi

Movie of a CUBIC cleared embryo stained with DAPI at Stage XX.2

A Stage XX.2 CUBIC cleared, DAPI stained embryo imaged with LSFM shown rotating along the longitudinal axis.

